# Single-cell transcriptome analysis illuminating the characteristics of species-specific innate immune responses against viral infections

**DOI:** 10.1101/2022.12.06.519403

**Authors:** Hirofumi Aso, Jumpei Ito, Haruka Ozaki, Yukie Kashima, Yutaka Suzuki, Yoshio Koyanagi, Kei Sato

**Affiliations:** Division of Systems Virology, Department of Microbiology and Immunology, The Institute of Medical Science, The University of Tokyo, Tokyo 1088639, Japan; Institute for Life and Medical Sciences, Kyoto University, Kyoto 6068507, Japan; Graduate School of Pharmaceutical Sciences, Kyoto University, Kyoto 6068501, Japan; Bioinformatics Laboratory, Faculty of Medicine, University of Tsukuba, Tsukuba 3050821, Japan; Center for Artificial Intelligence Research, University of Tsukuba, Tsukuba 3058577, Japan; Laboratory of Systems Genomics, Graduate School of Frontier Sciences, The University of Tokyo, Kashiwa 2778561, Japan; International Research Center for Infectious Diseases, The Institute of Medical Science, The University of Tokyo, Tokyo 1088639, Japan; International Vaccine Design Center, The Institute of Medical Science, The University of Tokyo, Tokyo 1088639, Japan; Graduate School of Medicine, The University of Tokyo, Tokyo 1130033, Japan; Graduate School of Frontier Sciences, The University of Tokyo, Kashiwa 2778561, Japan; Collaboration Unit for Infection, Joint Research Center for Human Retrovirus infection, Kumamoto University, Kumamoto, Japan; CREST, Japan Science and Technology Agency, Kawaguchi 3320012, Japan

**Keywords:** innate immunity, mammal, virus infection, single-cell RNA-sequencing, tensor

## Abstract

Bats harbor various viruses without severe symptoms and act as their natural reservoirs. The tolerance of bats against viral infections is assumed to originate from the uniqueness of their immune system. However, how immune responses vary between primates and bats remains unclear. Here, we characterized differences in the immune responses by peripheral blood mononuclear cells to various pathogenic stimuli between primates (humans, chimpanzees, and macaques) and bats (Egyptian fruit bats) using single-cell RNA sequencing. We show that the induction patterns of key cytosolic DNA/RNA sensors and antiviral genes differed between primates and bats. A novel subset of monocytes induced by pathogenic stimuli specifically in bats was identified. Furthermore, bats robustly respond to DNA virus infection even though major DNA sensors are dampened in bats. Overall, our data suggest that immune responses are substantially different between primates and bats, presumably underlying the difference in viral pathogenicity among the mammalian species tested.

## Introduction

Although a virus can infect various animal species, the pathogenicity of the infection can differ among host species. For example, Old World monkeys, including rhesus macaques (*Macaca mulatta*), are naturally infected with Cercopithecine herpesvirus 1 (also known as B virus) without any observable disorders, while humans (*Homo sapiens*) exhibit severe disorders after infection^1^. Bat species are naturally infected with a variety of viruses and behave as natural reservoirs of human pathogenic viruses^2^. For example, Marburg virus infection causes severe symptoms in humans but not in Egyptian fruit bats (*Rousettus aegyptiacus*), a putative natural host of this virus^3^. One possible factor that could define the differences in viral pathogenicity among host species is the difference in innate immune responses. For example, a previous study reported that Egyptian fruit bats lack the induction of proinflammatory cytokines, including *CCL8, FAS*, and *IL6*, which are related to disease severity in humans, upon Marburg virus infection, suggesting that the lack of cytokine induction is one of the reasons why Egyptian fruit bats exhibit asymptomatic infection with Marburg virus^4^.

Pathogen sensing is the initial step in triggering innate immune signaling. In a broad range of animals, including vertebrates, pathogen-associated molecular patterns (PAMPs) are recognized by pattern recognition receptors (PRRs) to induce subsequent immune responses^5-8^. In humans and mice (*Mus musculus*), double-stranded RNAs (dsRNAs), a PAMP for RNA viruses, are recognized by RNA sensors, such as RIG-I, MDA5, LGP2, TLR3, and TLR7/8^5,6^. Cytosolic DNAs, a PAMP for DNA viruses, are recognized by DNA sensors, such as cGAS, AIM2, IFI16, and TLR9^5,6,9^. Lipopolysaccharide (LPS), a PAMP for bacteria, is recognized by TLR4^5,6,10^. Once PAMPs are recognized by PRRs, type I interferons (IFNs) are produced, leading to the induction of IFN-stimulated genes (ISGs), which include many antiviral genes^5,6^.

In contrast to the similarities in the immune system between humans and mice, the immune system of bats is assumed to be quite different from that of humans in various aspects^11-13^. Genome analysis of Egyptian fruit bats showed expansion and diversification of immune-related genes, including type I IFN genes^14^. Transcriptome analysis showed that type I IFNs in the Australian black flying fox (*Pteropus alecto*) are constitutively expressed in unstimulated tissues, leading to the constitutive expression of ISGs^15^. These observations suggest that immunity in bats may be stronger than that in other mammals. In contrast, some studies have proposed that immune responses in bats are dampened, resulting in bats exhibiting stronger tolerance to various viruses^12,14,16^. In particular, it is known that critical molecules involved in viral DNA sensing, such as cGAS, AIM2, and IFI16, are dampened or genetically lost in some bat species, including Egyptian fruit bats^16,17^. These differences in innate immunity between humans and bats could be one of the reasons why viral pathogenicity differs between these two mammals.

Previous works have highlighted the uniqueness of the bat immune system using genomic analysis^14,15,17^, transcriptome analysis^4,18-20^, and molecular biological experiments that reconstituted a part of the bat immune system in cell culture systems^16,21,22^. However, it remains unclear how and to what extent the innate immune response to pathogenic stimuli varies among mammals. Particularly, it is unclear how different innate immune responses are elicited by viral infections in different cell types in each mammal. Here, we used peripheral blood mononuclear cells (PBMCs) from four mammalian species and three pathogenic stimuli and conducted single-cell RNA sequencing (scRNA-seq) analysis to elucidate the differences in innate immune responses against pathogenic stimuli.

## Results

### Experimental design

To illuminate the differences in immune responses to infectious pathogens among mammalian species, we isolated PBMCs from four mammals including humans (*Homo sapiens*, Hs), chimpanzees (*Pan troglodytes*, Pt), rhesus macaques (*Macaca mulatta*, Mm), and Egyptian fruit bats (*Rousettus aegyptiacus*, Ra) (**Fig. 1A**). These PBMCs were inoculated with herpes simplex virus type 1 (HSV-1; a DNA virus), Sendai virus (SeV; an RNA virus), or lipopolysaccharide (LPS; a proxy for bacterial infection). We verified that these PBMCs could be infected with and/or respond to these viruses and LPS stimulation by quantifying viral RNAs and the upregulation of proinflammatory cytokines (e.g., IL1B and IL6), ISGs (e.g., EIF2AK2 and DDX58) and IFNB1 (**Fig. S1A–C**).

**Figure 1.**
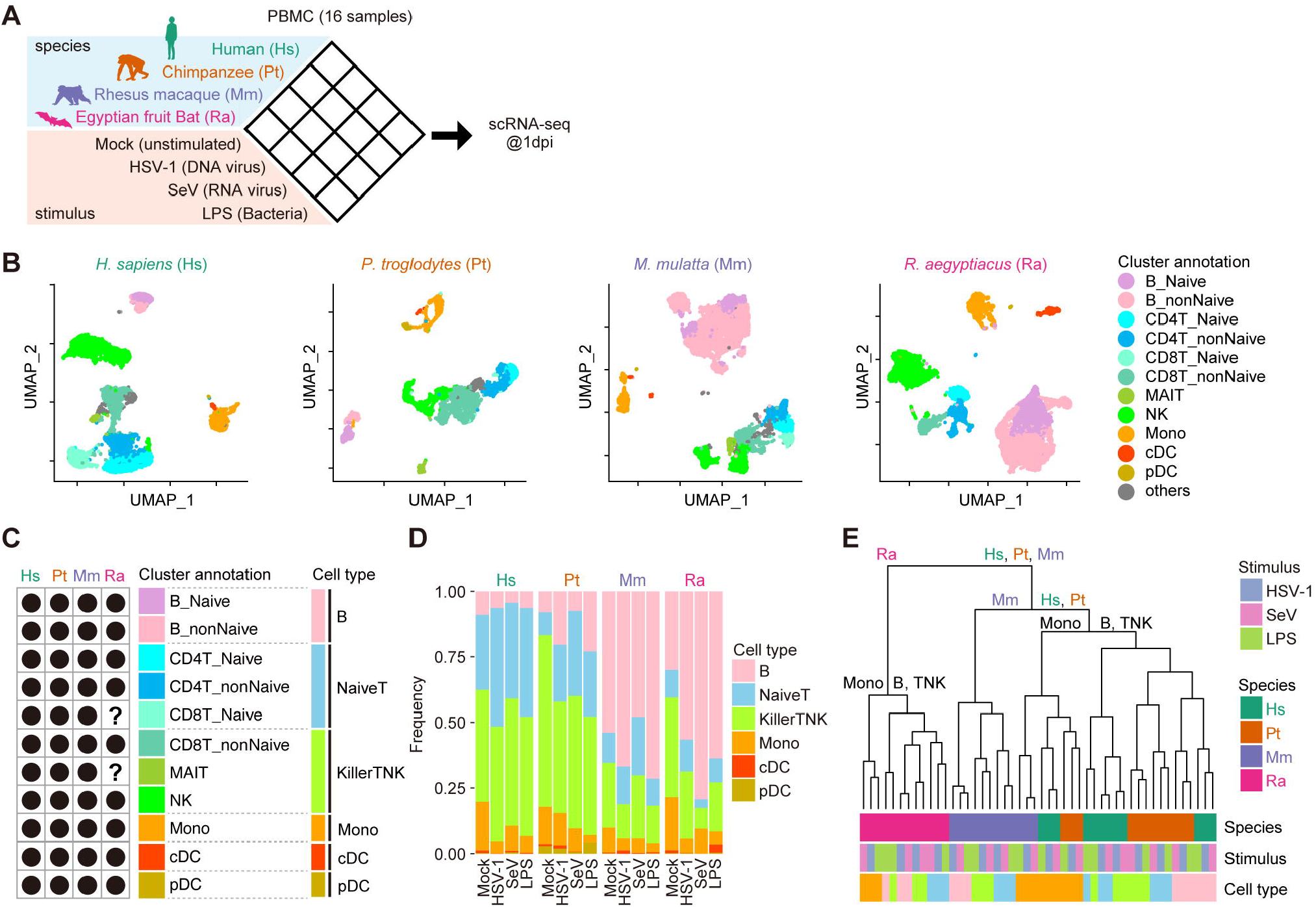
scRNA-seq analysis of PBMCs from four animal species inoculated with pathogenic stimuli. (A) Schematic of the experimental design. See also **Fig. S1**. (B) Uniform manifold approximation and projection (UMAP) plots representing the gene expression patterns of the cells from the four species. Each dot is colored according to the cell type. Gray dots indicate cells unassigned into any cell type. See also **Fig. S2**. (C) Comparison of identified cell types among the species. Dot: detected, question mark: undetected. The definitions of six species-common cell types are shown on the right side. See also **Fig. S2H**. (D) The cellular compositions of PBMC samples. The compositions according to the six common cell types are shown. (E) Hierarchical clustering analysis of 48 pseudobulk datapoints (4 animal species x 3 stimuli x 4 cell types = 48 conditions) based on PC1-30 calculated from the fold-change values (respective stimulus versus unstimulated) for gene expression.

To analyze immune responses to stimuli at single-cell resolution, we performed scRNA-seq analysis of 16 types of PBMC samples: four mammalian species (Hs, Pt, Mm, and Ra) versus four conditions (mock infection/stimulation, HSV-1 infection, SeV infection, and LPS stimulation) using the 10x Genomics Chromium platform. After filtering low-quality cells, a total of 40,717 cells from the 16 samples were used in the following analysis.

### The cellular composition of PBMCs from primates and bats

We characterized the cellular composition of PBMCs from each mammalian species by annotating the cell type of individual single cells. To establish a common classification system for the cells from the different mammalian species, we first identified cell types present in multiple species (**Fig. 1B and 1C**). As cell types detected in multiple species, naïve B cells, non-naïve B cells (including memory B cells and intermediate B cells), naïve CD4+ T cells, non-naïve CD4+ T cells (including central memory CD4+ T cells, effector memory CD4+ T cells, proliferating CD4+ T cells, and regulatory T cells), naïve CD8+ T cells, non-naïve CD8+ T cells (including central memory CD8+ T cells, effector memory CD8+ T cells, and proliferating CD8+ T cells), natural killer (NK) cells, mucosal-associated invariant T cells (MAITs), monocytes (Monos), conventional dendritic cells (cDCs), and plasmacytoid DCs (pDCs) were identified (**Fig. 1C**). Known marker genes for each cell type in humans were detected in the corresponding cell type in the unstimulated samples from the other animal species (**Fig. S2G**). Although most cell types were detected in all four species investigated, naïve CD8+ T cells and MAITs were undetectable in bat PBMCs, presumably because the cell numbers of these populations were relatively low in bats and/or the transcriptomic signatures of naïve CD4+ T cells and non-naïve CD8+ T cells were too similar in bats (hereafter we simply referred to Egyptian fruit bats as “bats”) (**Fig. 1C**). This result was consistent with a previous study, in which clear clusters of naïve CD8+ T cells and MAITs were not detected^23^. To establish a cellular classification system for the comparative transcriptome analysis, we defined six species-common cell types, namely, B cells, naïve T cells, killer TNK cells, Monos, cDCs, and pDCs, according to similarities in expression patterns (**Fig. S2H**).

The composition of the six cell types exhibited different changes upon exposure to the stimuli in the different species (**Fig. 1D**). The frequency of monocytes decreased after stimulation in all four species, whereas the frequency of B cells changed differently among the animal species and stimuli. After SeV infection, the frequency of B cells was decreased in all four species. On the other hand, after HSV-1 infection, the frequency of B cells was decreased in only humans.

### The differences in the immune response are large among animal species

To describe the differences in immune responses to various stimuli in specific cell types among animal species, we first calculated the average expression levels of appropriate genes in each condition (4 animal species × 4 stimuli × 6 cell types = 96 conditions). Using this “pseudobulk” transcriptome dataset, we first investigated which axis (i.e., animal species, stimulus, and cell type) was the most impactful element in shaping the expression patterns of immune cells. Thereby, we calculated the fold-change (FC) values of gene expression levels between unstimulated and corresponding stimulated conditions and performed principal component analysis (PCA) on the FC values. Subsequently, hierarchical clustering analysis was performed according to principal components (PCs) 1-30. The transcriptome data were first branched according to the animal species and then branched according to the cell type followed by the stimulus (**Fig. 1E**). This suggested that the difference in host species was the more impactful element in shaping the immune system, having a greater impact than the type of stimulus and cell type. In particular, our dataset showed that bat PBMCs exhibited different transcriptomic patterns irrespective of the type of stimulus and cell type compared to the PBMCs from the other three species used. Our results suggest that bats respond to pathogens in a different manner than primates.

### Extraction of species-specific immune responses

We next characterized the differences in the immune responses to pathogenic stimuli among animal species. The FC values of our pseudobulk transcriptome dataset were represented by a four-mode tensor (4 animal species × 3 stimuli × 6 cell types × 7557 orthologous genes). To characterize this extraordinary high-dimensionality transcriptome dataset, we utilized Tucker decomposition, a method of tensor decomposition (**Fig. 2A**). In this analysis, we excluded cDC and pDC data due to many missing values. Tucker decomposition generated a core tensor and four-factor matrices (A1–A4) related to the four axes (animal species, stimulus, cell type, and gene). For example, the factor matrix A1 (for host species) included three latent factors (L1_1, L1_2, and L1_3), which could be regarded to represent common, bat-specific, and macaque-specific expression patterns, respectively (**Fig. 2B**).

**Figure 2.**
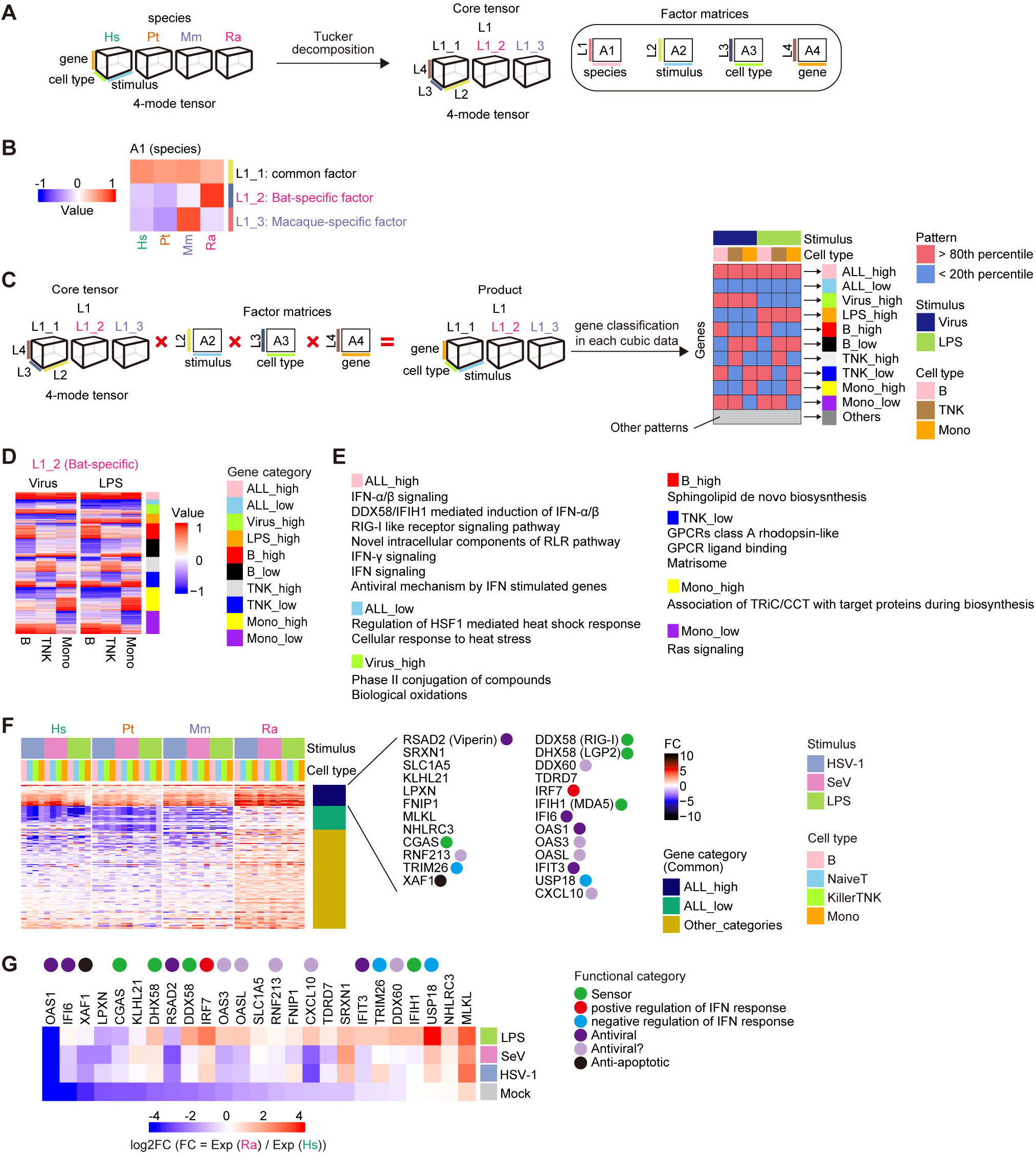
Characterization of species-specific immune responses using a tensor decomposition framework. (A) Tensor decomposition of the fold-change values for pseudobulk transcriptome data. (B) Heatmap representing a latent factor matrix relating to species. Columns indicate the animal species, and rows indicate the latent factors representing species-common (L1_1), bat-specific (L1_2), and macaque-specific (L1_3) factors. See also **Fig. S3A–B**. (C) Classification of genes according to the differential patterns of the latent factors related to species. For each of the species-common (L1_1), bat-specific (L1_2), and macaque-specific (L1_3) factors, the product of the core tensor and three latent factor matrices related to stimulus, cell type, and gene was calculated (left), and the genes were classified into 11 categories according to the binary patterns for each calculated product (right). See also **Fig. S3C–F**. (D) Heatmap representing the values of the products calculated in **Figure 2C**. From the three products, the data related to the bat-specific factor (L1_2) are shown. Each row indicates the respective gene. The color keys shown on the right of the heatmap indicate gene categories. See also **Fig. S3G–L**. (E) GO terms enriched in each gene category relating to the bat-specific factor. GO terms with a false discovery rate (FDR) <= 0.1 and an odds ratio >= 1 are shown. (F) Heatmap representing the induction levels of ALL_high genes for the bat-specific factor. Additional classification according to the gene classification of the species-common factors is shown to the right of the heatmap. Genes categorized as ALL_high in both the species-common factor and the bat-specific factor are shown on the right side. The colored circle indicates the functional category of the gene. (G) Heatmap representing the relative expression levels (bats versus humans) of the genes shown in **Figure 2F**.

To characterize species-specific immune responses, we developed a gene classification system according to the pattern of the species-associated latent factor in the tensor decomposition framework. First, we calculated the product of a core tensor and the three-factor matrices A2 (for stimulus), A3 (for cell type), and A4 (for gene) (**Fig. 2C and Fig. S3A–B**). Consequently, we obtained three cubic datasets with three axes, stimulus, cell type, and gene. These cubic data were related to L1_1 (for the common factor), L1_2 (for the bat-specific factor), or L1_3 (for the macaque-specific factor). Subsequently, we classified the genes into 10 categories according to their expression patterns in each cubic dataset (the results for the bat-specific (L1_2) and other factors (L1_1 and L1_3) are shown in **Fig. 2D, Fig. S3G, and Fig. S3I**, respectively). In the factor matrix A2 (for stimulus), the values for the latent factors related to HSV-1 and SeV were similar (**Fig. S3A**). Therefore, these two categories were integrated into the category “Virus” in the gene classification. Additionally, two cell type categories, NaiveT and KillerTNK, were integrated into the category “TNK” (**Fig. S3B**). The pattern for raw FC values supported that the gene classification by the tensor decomposition framework succeeded in extracting the characteristic patterns of gene expression alterations upon pathogenic stimuli (**Fig. S3J–L**).

### Differential dynamics of pathogen sensing and immune responses

To highlight the uniqueness of immunity in bats compared to that in primates, we focused on the expression pattern represented by the bat-specific factor (L1_2) and performed Gene Ontology (GO) analysis on the 10 gene categories (**Fig. 2E**). In the gene category “ALL_high”, which included genes upregulated particularly in bats regardless of the stimulus and cell type, GO terms related to innate immune responses, such as IFN signaling, DDX58/IFIH1-mediated induction of IFN, RIG-I like receptors (RLRs) signaling pathways, and the antiviral mechanism by ISGs, were enriched.

To dissect the “ALL_high” genes in the bat-specific factor, we further extracted the genes that belonged not only to the “ALL_high” category in the bat-specific factor but also to that in the common factor (L1_1). This fraction represented genes that were upregulated by stimuli in all species but whose induction levels were highest in bats. These genes included various PPRs, such as RIG-I-like receptors (RLRs) (RIG-I, LGP2, and MDA5) and cGAS, a DNA sensor, suggesting that these genes were upregulated to higher levels in bats than in the other species across the cell types and stimuli (**Fig. 2F**). These higher FC values in bats could be explained by two possibilities. First, the expression levels of these genes after stimulation were higher in bats than in primates. Second, the basal expression levels of these genes in bats were lower than those in primates. Therefore, we calculated the relative expression levels of these genes in bats compared to humans and showed that the basal expression levels of these genes were lower in bats than in humans (**Fig. 2G**). These results suggest that the induction dynamics of these PRRs in bats are likely different from those in primates, possibly leading to the differences in the induction of immune responses.

### Robust immune responses to a DNA virus in bats

As critical DNA sensors, such as cGAS, AIM2, IFI16, and TLR9, are dampened or genetically lost in bat species^16,17,24^, it has been hypothesized that bats, including Egyptian fruit bats, cannot efficiently activate innate immune responses against DNA viruses. To test this hypothesis, we analyzed the IFN response upon HSV-1 (a DNA virus) infection by analyzing the induced levels of “core^mamm^ ISGs”, a set of genes that are commonly induced by type I IFNs across mammals that were defined in a previous study ^25^. Intriguingly, we found that the core^mamm^ ISGs were upregulated upon HSV-1 infection in most cell types in bats (**Fig. 3A**). The induced levels were comparable to those induced by SeV (an RNA virus) infection and higher than those induced by LPS stimulation. Furthermore, the induced levels in bats were comparable to those in primates. This suggests that immune cells in bats can sense and respond to HSV-1 infection even though critical DNA sensors are dampened.

**Figure 3.**
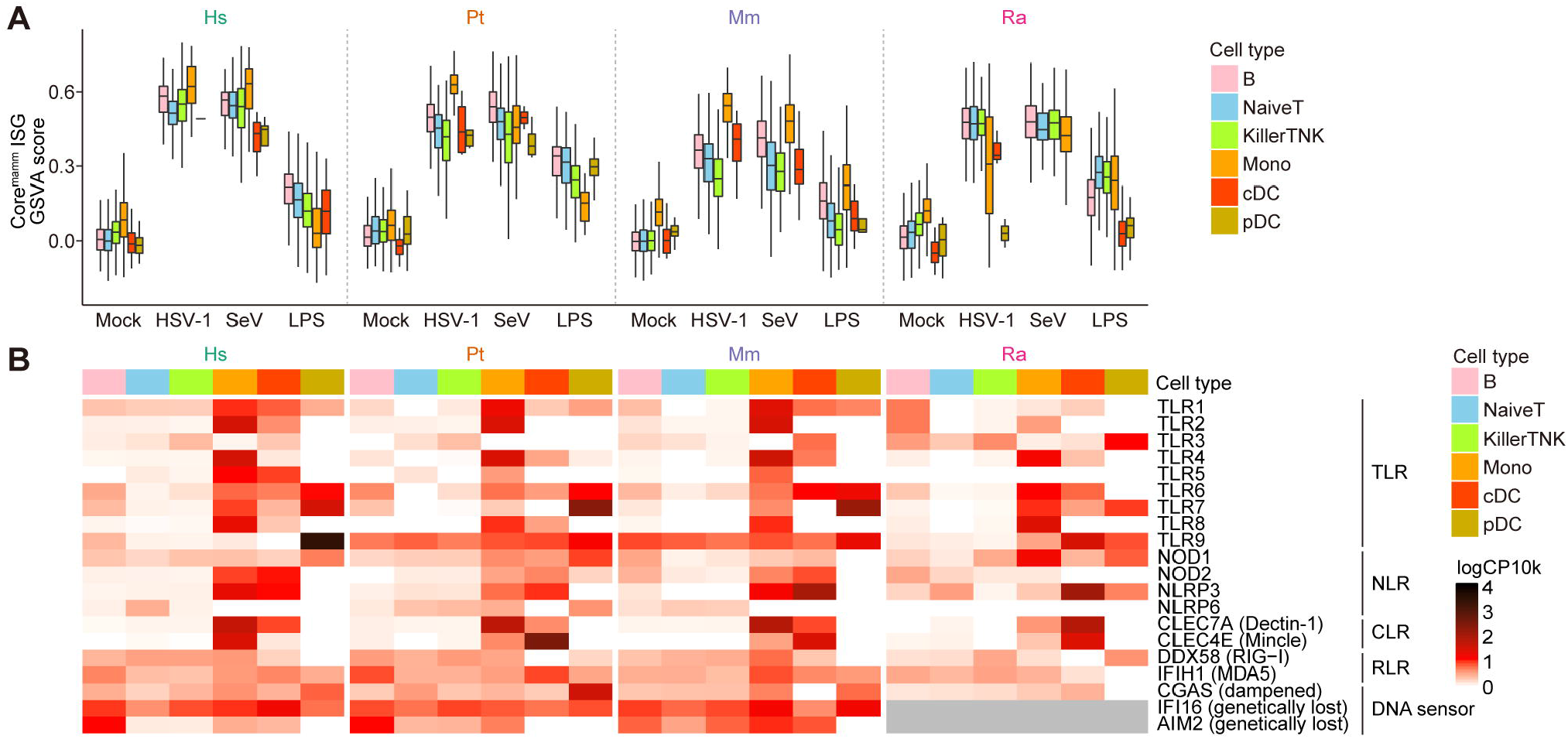
Robust immune responses to a DNA virus in bats. (A) Boxplot of the expression levels of core^mamm^ ISGs in every single cell. The Y-axis indicates the global expression level (GSVA score) of the core^mamm^ ISGs. (B) Heatmap representing the mean expression levels of sensor genes. The mean values were calculated without using the information for the stimulus.

To address the possibility that pathogen sensors other than DNA sensors contribute to the sensing of HSV-1 infection in bats, we examined the expression levels of various PRRs (**Fig. 3B**). The expression of some PRRs, including TLR3, a dsRNA sensor associated with HSV-1 sensing in humans and mice^26^, was detected not only in primates but also in bats, suggesting the possibility that these PRRs compensate in the response to HSV-1 infection in bats (see **Discussion**).

### Identification of bat-specific subsets of monocytes

Next, we investigated cellular subsets within the cell types that are characteristic in bats to explain the differences in immune responses among the species. We particularly searched for cellular subsets that specifically appeared after pathogenic stimulus exposure in each species according to the dimensionality reduction analysis of transcriptome data. In humans, chimpanzees, and macaques, no subset appeared in any cell type after stimulation (**Fig. S4A**). Similarly, such subsets were not identified in T/NK or B cells in bats. In contrast, we found that two subsets of bat monocytes (referred to as Clusters 5 and 7) specifically appeared after stimulation (**Fig. 4A**). To validate whether these subsets (Clusters 5 and 7) are unique in bats, we identified marker genes for these clusters and subsequently examined whether the marker genes were expressed in monocytes from the other animal species. The marker genes for Cluster 5 (referred to as C5 markers) were not highly expressed in any cluster of monocytes from primates (**Fig. 4B**). Furthermore, high expression levels of C5 markers in bat monocytes were found only after stimulation. This suggested that Cluster 5 was not only bat-specific but also specifically induced by pathogenic stimuli. Unlike the C5 markers, the marker genes for Cluster 7 (C7 markers) were highly expressed not only in bat Cluster 7 but also in some monocytes in primates (**Fig. 4C**). Although cells with higher expression of C7 markers were induced upon stimulation in both bats and primates, these cells in primates did not form a separate cluster similar to Cluster 7 in bats (**Fig. S4B**). Furthermore, the proportions of Clusters 5 and 7 differed depending on the stimulus: HSV-1-infected and LPS-stimulated samples showed the highest frequencies of Clusters 5 and 7, respectively (**Fig. 4D**).

**Figure 4.**
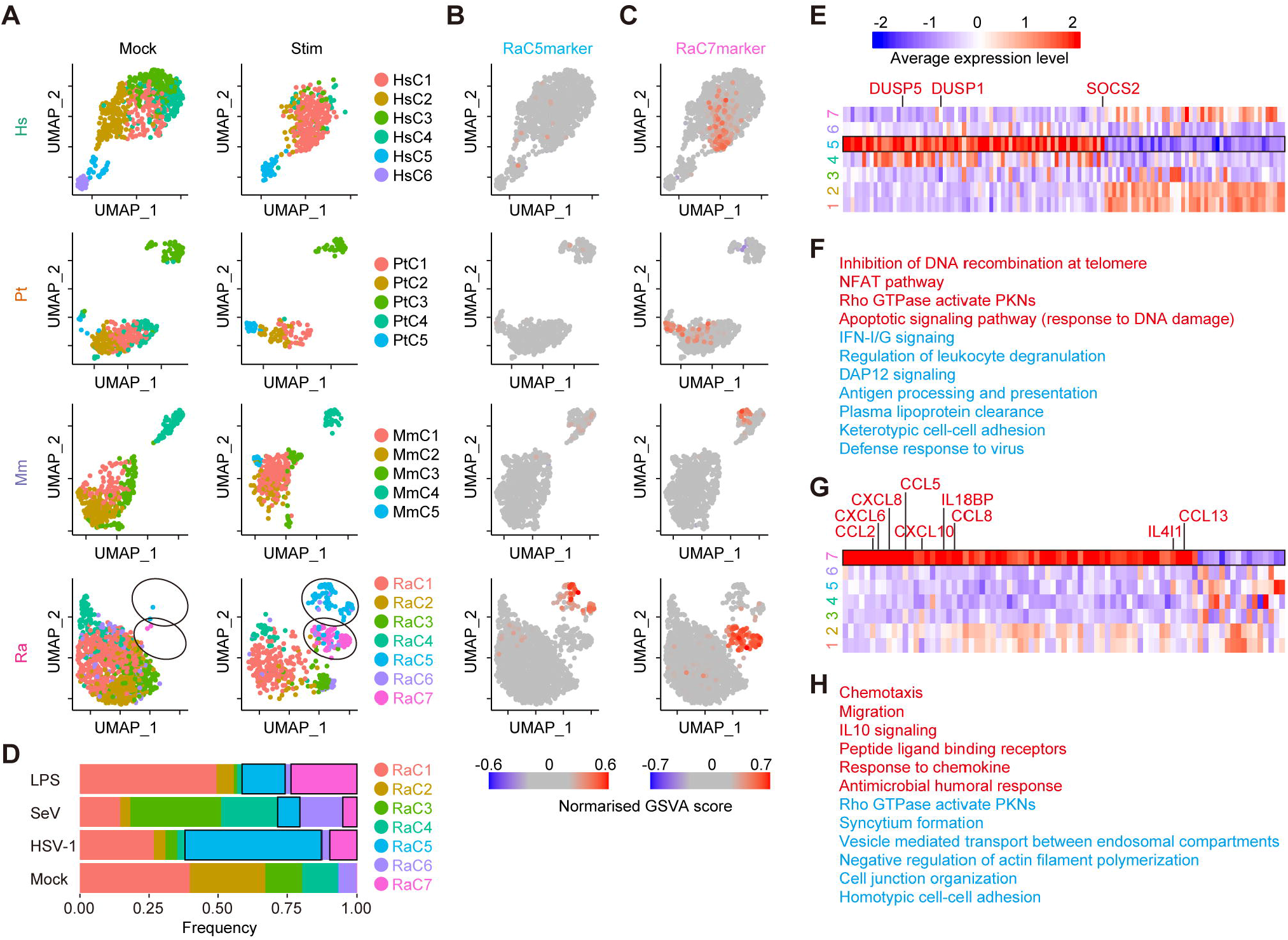
Identification of bat-specific subsets of monocytes. (A) UMAP plots representing the gene expression patterns of monocytes from the four species. The dots are colored according to the cell cluster defined for each animal species. See also **Fig. S4A**. (B, C) UMAP plots representing the average expression levels of marker genes for Cluster 5 [C5markers] (B) and Cluster 7 [C7markers] (C). See also **Fig. S4B**. (D) The cellular composition of bat monocytes. The composition is shown according to the cluster. The black frame indicates Clusters 5 and 7 in stimulated samples. (E) Heatmap representing the mean expression levels of differentially expressed genes (DEGs) in Cluster 5 of bat monocytes. (F) Summary of the GO terms enriched in DEGs in Cluster 5. GO terms enriched in up- and downregulated genes are shown in red and blue, respectively. (G) Heatmap representing the mean expression levels of differentially expressed genes (DEGs) in Cluster 7 of bat monocytes. (H) Summary of the GO terms enriched in DEGs in Cluster 7. GO terms enriched in up- and downregulated genes are shown in red and blue, respectively.

To characterize these two clusters, we identified differentially expressed genes (DEGs) in Clusters 5 and 7 compared to the other clusters of bat monocytes. According to GO analysis, Cluster 5 was characterized by lower expression of ISGs (**Fig. 4E, 4F**). Additionally, Cluster 5 highly expressed known suppressors of the inflammatory response, such as DUSP1, DUSP5, and SOCS2^27-29^. On the other hand, Cluster 7 could be characterized by a higher expression of various cytokines related to chemotaxis (**Fig. 4G**), including CXCL6, IL18BP, CXCL8, CCL2, CCL8, CCL13, CCL5, CXCL10, IL15, and IL4I1 (https://www.gsea-msigdb.org/gsea/msigdb/human/geneset/GOBP_CELL_CHEMOTAXIS.html) (**Fig. 4G, 4H**). Overall, we established that there are two unique subsets of bat monocytes with different characteristics (see **Discussion**).

## Discussion

Differences in viral pathogenicity among host species are thought to be attributed to differences in immune responses against viral infections among the species^30^. However, it remains unclear how immune responses, particularly innate immunity against viral infections, differ among host species. In the present study, we performed scRNA-seq on 16 types of PBMC samples, derived from a combination of four host species and four infection conditions (**Fig. 1A**), and showed that the differences in the immune responses among the host species were more impactful than those among both the stimuli and the cell types (**Fig. 1E**). In particular, the transcriptomic changes after pathogenic stimulation in bats differed from those in primates. Furthermore, we established a bioinformatic pipeline to characterize species-specific immune responses from transcriptome profiles with extraordinarily high dimensions (4 animal species × 3 stimuli × 4 cell types × 7,557 orthologous genes) (**Fig. 2A**). Our study provides fundamental data to identify differences in innate immune systems among mammalian species that partly explain the differences in viral pathogenicity among host species.

It is known that two DNA sensing pathways mediated by STING^16^ and PYHIN proteins, including AIM2 and IFI16^17^, are dampened in bats, including Egyptian fruit bats. In addition, a previous study using a cell line derived from big brown bats (*Eptesicus fuscus*) suggested that the TLR9-mediated DNA sensing pathway is also weakened in bats^24^. Based on these observations, it was hypothesized that the ability to sense DNA virus infection is weakened in bats^12,13^. However, we showed that bat PBMCs robustly induced IFN responses upon infection with the DNA virus HSV-1 (**Fig. 3A**). This suggests that bats can initiate an innate immune response after infection with DNA viruses (at least HSV-1) and that bats have another pathway to sense DNA viruses. An alternative possibility is that the IFN response in response to HSV-1 infection was triggered by sensing viral molecules other than DNAs: it is known that, in humans and mice, dsRNA sensing by TLR3 plays an important role in responding to HSV-1 infection^26,31^. Furthermore, the Egyptian fruit bat genome encodes an intact TLR3 gene (NCBI Gene ID: 107510436), and bat immune cells express TLR3 (**Fig. 3B**). These data suggest that in bats, bat TLR3 may compensate for the immune responses induced by DNA sensors, leading to IFN responses to HSV-1 infection.

To characterize the bat-specific innate immune responses based on ultrahigh-dimensionality transcriptome data (4 animal species × 4 stimuli × 6 cell types × 7,557 orthologous genes), we established an analytical framework utilizing tensor deconvolution (**Fig. 2A**). This framework could i) extract a species-specific effect on gene expression changes, ii) compare the effects among the cell types and the stimuli, and iii) classify genes according to the differential pattern of a species-specific effect among the cell types and the stimuli. Using this framework, we found that the expression levels of key DNA and RNA sensors, including cGAS, RIG-I, MDA5, and LGP2, were highly induced in bats compared with primates, regardless of the cell type or stimulus (**Fig. 2F**). Furthermore, the basal expression levels of these PRRs in bats were lower than those in humans (**Fig. 2G**). On the other hand, after stimulation, the expression levels of these PRRs in bats were comparable to those in humans. These results suggest that the induction dynamics of these PRRs in bats are likely different from those in primates, leading to the differences in the induction of immune responses. Indeed, several antiviral ISGs, such as IFI6 and IFIT3, exhibited expression dynamics similar to those of these PRRs (**Fig. 2F, 2G**). These differences could be one of the reasons why immune responses differ between bats and primates.

Another factor that can explain the differences in immune responses among host species is the presence of species-specific cellular subsets. In bat monocytes, we identified two subsets that were specifically induced by stimuli (i.e., Clusters 5 and 7) (**Fig. 4A**). Cluster 5 was a bat-specific subset induced preferentially by HSV-1 infection (**Fig. 4B, 4D**). Interestingly, even though Cluster 5 was induced after stimulation, Cluster 5 exhibited lower expression of ISGs and higher expression of immunosuppressive genes (DUSP1, DUSP5, and SOCS2)^27-29^ (**Fig. 4E, 4F**). This observation suggests that the immune responses in Cluster 5 are downregulated presumably by negative feedback signaling and that Cluster 5 may contribute to controlling excessive immune activation in bats. On the other hand, Cluster 7 was identified as a monocyte subset that was mainly induced by LPS stimulation (**Fig. 4C, 4D**). Cluster 7 highly expressed several proinflammatory cytokines and chemokines (CXCL6, IL18BP, CXCL8, CCL2, CCL8, CCL13, CCL5, CXCL10, IL15, and IL4I1) (**Fig. 4G, 4H**). Cluster 7 may contribute to the recruitment of leukocytes since these cytokines are associated with the chemotaxis of neutrophils (CCL8, CXCL6, and CXCL8), basophils (CXCL8, CCL2, CCL5, CCL8, and CCL13), eosinophils (CCL5, CCL8, and CCL13), monocytes (CCL5, CCL8, and CCL13), T cells (CCL5, CCL8, CCL13, CXCL8, and CXCL10), and NK cells (CCL5 and CCL8) in humans and mice^32^ (https://docs.abcam.com/pdf/immunology/chemokines_poster.pdf). Based on the expression pattern of the marker genes for Cluster 7 (**Fig. 4C, S4B**), cellular subsets corresponding to Cluster 7 were also present in primate monocytes. However, these primate cells did not form a separate cluster in the dimensionality reduction analysis based on the transcriptome profile (**Fig. 4A**). These results suggest that the monocyte subset represented by Cluster 7 exhibits unique gene expression and thus may exert unique functions in bats. Although the specific functions of these monocyte subsets (Clusters 5 and 7) in immune responses in bats are still unclear, these unique subsets may contribute to bat-specific host immune responses.

## Limitations of the study

In the present study, we elucidated differences in innate immune responses among host species from various aspects. However, we did not address differences in the outcomes of the innate immune responses, such as differences in viral pathogenicity. Another limitation is that the bioinformatic resources we used, such as gene annotation, gene ontology, and cellular annotation, have been developed in a human-centric way. Therefore, there is the possibility that immune responses induced by species-specific genes and cell types were overlooked. Despite these limitations, we present valuable resources to illuminate differences in immune responses among host species, including Egyptian fruit bats, and clues to elucidate differences in viral pathogenicity among species. Further study to elucidate the functional consequences of these differences is needed to reveal the mechanisms by which bats can tolerate infections with various viruses.

## Acknowledgements

We would like to thank Naoko Misawa, Akiko Oide, Mai Suganami, and Kazumi Abe (The University of Tokyo), for technical support, Hiroo Imai (Kyoto University) for providing primate PBMCs, Ayuko Morita (Kyoto City Institute of Health and Environmental Sciences) for providing bat PBMCs, Yasushi Kawaguchi (The University of Tokyo) for providing HSV-1, and Takashi Irie (Hiroshima University) for providing SeV, Human Genome Center (the Institute of Medical Science, the University of Tokyo) for providing the super-computing resource SHIROKANE (http://sc.hgc.jp/shirokane.html).

This study was supported in part by AMED SCARDA Japan Initiative for World-leading Vaccine Research and Development Centers “UTOPIA” (JP223fa627001, to K.S.), AMED SCARDA Program on R&D of new generation vaccine including new modality application (JP223fa727002, to K.S.); AMED Research Program on Emerging and Re-emerging Infectious Diseases (JP22fk0108146, to Y.Kashima and K.S.; JP21fk0108494 to K.S.; 21fk0108425, to K.S.; 21fk0108432, to K.S.); AMED Research Program on HIV/AIDS (JP22fk0410039, to K.S.); JST PRESTO (JPMJPR22R1, to J.I.); AMED Moonshot Research and Development Program (JP21zf0127005, to H.O.); JST CREST (JPMJCR20H4, to K.S.); JSPS KAKENHI Grant-in-Aid for Early-Career Scientists (20K15767, to J.I.; 19K20394, to H.O.); JSPS Core-to-Core Program (A. Advanced Research Networks) (JPJSCCA20190008, to K.S.); JSPS Research Fellow DC1 (20J23299, to H.A.).

## Author Contributions

H.A. mainly performed bioinformatics analysis. J.I. and H.O. supervised the bioinformatics analysis. Y.Kashima mainly performed the experiments. Y.S., Y.Koyanagi, and K.S. supervised the experiments. K.S. and Y.Koyanagi provided reagents. K.S. conceived and designed the experiments. H.A. and J.I. wrote the initial manuscript. All authors reviewed and edited the manuscript.

## STAR Methods

### Resource Availability

#### Lead contact

Further information and requests for resources and reagents should be directed to and will be fulfilled by the Lead Contact, Kei Sato.

#### Materials availability

All unique reagents generated in this study are listed in the Key Resources table and available from the Lead Contact with a completed Materials Transfer Agreement.

#### Data and code availability

- Single-cell RNA-seq data have been deposited in the GEO database (GSE218199) and are publicly available as of the date of publication. Original data to describe figures in this paper have been deposited at Mendeley (DOI: 10.17632/kg3dfkyjv5.1) and are publicly available as of the date of publication.
- All original code has been deposited at GitHub (https://github.com/TheSatoLab/scRNA-seq_PBMC_Animals_Aso_et_al) and is publicly available as of the date of publication.
- Any additional information required to reanalyze the data reported in this paper is available from the Lead Contact upon request.

### Experimental Model and Subject Details

#### Ethics Statement

All protocols involving specimens from animals were performed in accordance with the Science Council of Japan’s Guidelines for the Proper Conduct of Animal Experiments. The protocols were approved by the Institutional Animal Care and Use Committee of Kyoto University (approval IDs: 2017-B-5, 2019-C-9, and 2020-C-5). All protocols involving specimens from humans recruited at Kyoto University were reviewed and approved by the Institutional Review Boards of Kyoto University (approval ID: G1089). All human subjects provided written informed consent. All protocols for the use of human specimens were reviewed and approved by the Institutional Review Boards of The Institute of Medical Science, The University of Tokyo (approval ID: 2019-55) and Kyoto University (approval ID: G1089).

#### Cells

Vero cells (obtained from the Laboratory of Bernard Roizman, University of Chicago, USA)

LLC-MK2 cells (rhesus macaque kidney epithelial cells) (CCL-7, ATCC)

#### PBMC collection

Human peripheral blood was obtained from the arm vein. To obtain chimpanzee peripheral blood, a chimpanzee was anesthetized for a regular health examination. Anesthesia was induced with intramuscular administration of the combination of 0.012 mg/kg medetomidine (Meiji Seika Pharma Co., Ltd.), 0.12 mg/kg midazolam (Sand Co., Ltd.), and 3.5 mg/kg ketamine and maintained with constant rate infusion (4-10 mg/kg/h) of propofol (1% Diprivan, Sand Co., Ltd.). Peripheral blood was obtained from the femoral vein. To obtain rhesus macaque peripheral blood, a rhesus macaque was anesthetized. Anesthesia was induced with intramuscular administration of 8 mg/kg ketamine (Fujita Pharm, Tokyo) followed by deep anesthetization using an intravenous injection of sodium pentobarbital (30 mg/kg). Peripheral blood was obtained by cardiac puncture before exsanguination and perfusion. Bat peripheral blood was obtained from the cephalic vein in the patagium. PBMCs were isolated from peripheral blood by density gradient centrifugation using Ficoll-Paque™ Plus (Cytiva, Cat# 17144003).

#### HSV-1 preparation and titration

HSV-1 (strain F; GenBank accession number: GU734771)^33^ was prepared as previously described^26^ and kindly provided by Dr. Yasushi Kawaguchi (The Institute of Medical Science, The University of Toyo, Japan). To titrate viral infectivity, prepared virus was diluted 10-fold in Medium 199 (Thermo Fisher Scientific, Cat# 11825015) containing 1% FCS, and Vero cells were infected with dilutions of the virus at 37 °C. At one hour postinfection, the culture medium was replaced with Medium 199 containing 160 μg/ml human γ-globulin (Sigma Aldrich, G4386-25G), and the cells were cultured at 37 °C for 2–3 days. To calculate the viral titer [plaque forming unit (PFU)], the number of plaques per well was counted.

#### SeV preparation and titration

SeV (strain Cantrell, clone cCdi; GenBank accession number: AB855654) was prepared as previously described^34^ and kindly provided by Dr. Takashi Irie (Hiroshima University, Japan). To titrate viral infectivity, prepared virus was diluted 10-fold in Dulbecco’s modified Eagle’s medium (DMEM) (Sigma□Aldrich, Cat# D6046-500ML) containing 10% FCS, and LLC-MK2 cells were infected with dilutions of the virus at 37 °C. At one hour postinfection, the cells were washed with PBS and cultured with DMEM containing 10% FCS at 37 °C. At one day postinfection, the infected cells were fixed with acetone/methanol. To calculate the viral titer [cell infectious unit (CIU)], the fixed cells were stained with a rabbit anti-SeV polyclonal antibody^35^ as the primary antibody and an Alexa 488-conjugated goat anti-rabbit IgG antibody (Thermo Fisher Scientific, Cat# A-11008) as the secondary antibody, and the number of fluorescent foci per well was counted.

### Method Details

#### Infection and stimulation

One million PBMCs were maintained in 500 μl RPMI 1640 medium (Sigma□Aldrich, Cat# R8758-500ML) and infected with HSV-1 or SeV at a multiplicity of infection of 0.1. To mimic microbial infection, LPS (Sigma□Aldrich, Cat# L5024-10MG) was added at a final concentration of 200 ng/ml. At one day post infection, infected/stimulated PBMCs were centrifuged, resuspended in PBS, and used for bulk RT–qPCR and scRNA-seq (see below).

#### RT–qPCR

RT–qPCR was performed as previously described^36^. Briefly, cellular RNA was extracted using the QIAamp RNA Blood Mini Kit (Qiagen, Cat# 52304) and then treated with an RNase-free DNase set (Qiagen, Cat# 79254). cDNA was synthesized using SuperScript III reverse transcriptase (Thermo Fisher Scientific, Cat# 18080044) and random primers (Thermo Fisher Scientific, Cat# 48190011). RT–qPCR was performed using Power SYBR Green PCR Master Mix (Thermo Fisher Scientific, Cat# 4367659) and the primers listed in **Table S1**. For RT–qPCR, the CFX Connect Real-Time PCR Detection System (Bio-Rad) was used.

#### Sequencing of scRNA-seq libraries

scRNA-seq libraries were constructed using the Chromium Next GEM Single Cell 3’ Kit according to the manufacturer’s instructions (10x Genomics). Briefly, cells, gel beads, and oil were loaded onto the Chromium platform to generate single-cell gel beads-in-emulsion (GEMs). Barcoded cDNAs were pooled for amplification, and adaptors and indices for sequencing were added. The evaluation was conducted using a BioAnalyzer (Agilent Technologies). The libraries were sequenced with paired-end reads using the NovaSeq6000 platform (Illumina).

#### Genome sequence dataset

Genome sequences of the animal species including humans (GRCh38.p13, RefSeq accession: GCF_000001405.39), chimpanzees (Clint_PTRv2, RefSeq accession: GCF_002880755.1), rhesus macaques (Mmul_10, RefSeq accession: GCF_003339765.1), and Egyptian fruit bats (mRouAeg1.p, RefSeq accession: GCF_014176215.1) were obtained from NCBI RefSeq (www.ncbi.nlm.nih.gov/genome). From the genome sequences, ALT contig sequences were excluded. The genome sequences of viruses including HSV-1 (strain: F, accession: GU734771.1) and SeV (strain: Cantell clone cCdi, accession: AB855654.1) were also obtained from NCBI RefSeq. A custom reference genome sequence for each animal species was generated by adding the genome sequences of HSV-1 and SeV to the genome sequence of the animal species.

#### Gene annotation and ortholog information

Gene annotations of humans (GRCh38.p13, Release 109.20200228), chimpanzees (Clint_PTRv2, Release 105), rhesus macaques (Mmul_10, Release 103), and Egyptian fruit bats (mRouAeg1.p, Release 101) were obtained from NCBI RefSeq. From the gene annotations, only the records for protein_coding, transcribed_pseudogene, lncRNA, pseudogene, antisense_RNA, ncRNA_pseudogene, V_segment, V_segment_pseudogene, C_region, C_region_pseudogene, J_segment, J_segment_pseudogene, and D_segment were extracted according to the CellRanger tutorial (https://support.10xgenomics.com/single-cell-gene-expression/software/pipelines/latest/using/tutorial_mr). In addition, to quantify viral RNA abundance, the records for viruses were added. The whole viral genome was treated as a single exon, and a total of four lines (the positive and negative strands of HSV-1 and SeV) were added.

A list of orthologous genes between humans and the other animal species (chimpanzees, rhesus macaques, and Egyptian fruit bats) was obtained from NCBI on July 26th, 2021 (https://ftp.ncbi.nih.gov/gene/DATA/gene_orthologs.gz). From the file, the records for orthologs between humans (taxonomy ID: 9606) and chimpanzees (taxonomy ID: 9598), rhesus macaques (taxonomy ID: 9544), or Egyptian fruit bats (taxonomy ID: 9407) were extracted.

The ortholog list from NCBI lacked information on some critical immune-related genes of Egyptian fruit bats, such as CD4 and IRF1. Therefore, we retrieved information from the Bat1K gene annotation^37^ (https://bat1k.com): First, we made a custom gene annotation for Egyptian fruit bats by adding information from the Bat1K gene annotation to the RefSeq gene annotation. Second, we extracted exons in the Bat1K gene annotation that overlapped with exons in the RefSeq gene annotation by using the bedtools intersect command with the wao option (v2.30.0)^38^. In this step, the exons in the Bat1K gene annotation that did not overlap with the exons in the RefSeq gene annotation were also extracted and added to custom gene annotations as additional genes.

Next, the exons that contained overlaps and had the same gene name (the same symbol or known to be an ortholog) were added to custom gene annotations as an alternative splicing variant of the gene. Then, the remaining overlapping exons were processed by determining which information (RefSeq or Bat1K) should be used preferentially. The criteria were as follows: i) genes whose symbols are not prefixed with “LOC” were given priority, ii) genes whose symbols are included in the human gene list were given priority, and iii) information from RefSeq was given priority otherwise. According to these criteria, the annotation with the higher priority (RefSeq or Bat1K) was selected and used in the custom gene annotation.

As a result of the integration of gene annotations, the number of orthologous genes in the custom gene annotation of bats increased from 16374 to 16903. Importantly, immune-related genes that were not defined in the RefSeq gene annotation, such as TLR1, IRF1, and CD4, were added to the custom gene annotation.

Considering the orthologous relationships, we prepared three types of gene sets for each animal species: i) “all genes”, including all genes in the animal species; ii) “genes shared with humans”, including genes with orthologs in humans; and iii) “common genes”, genes shared among the four analyzed animal species. Unless otherwise noted, “all genes” were used up to cell annotation, and “common genes” were used after cell annotation.

#### Processing scRNA-seq data for generating count matrices

Gene expression count matrices for scRNA-Seq data were generated using CellRanger (v6.0.1) (10x Genomics). First, we built a custom reference for each animal species from the custom reference genome sequence and custom gene annotation using the “cellranger mkref” command. Subsequently, we generated unique molecular identifier (UMI)-based count matrices from the raw scRNA-seq data and custom references using the “cellranger count” command with default settings.

#### Quality control (QC) of scRNA-seq data

First, we removed cells with abnormal genes per cell (genes/cell) and counts per cell (counts/cell) values using the Seurat package (v4.0.4)^39^: Cells with 800–5,000 genes/cell or 1,200–25,000 counts/cell were extracted. Second, we removed nontargeted cells in the present study. We annotated the cell type of individual cells using Azimuth (v0.4.3), a reference-based cell annotation prediction program (https://azimuth.hubmapconsortium.org), and cells annotated as erythrocytes, hematopoietic stem cells, innate lymphoid cells, and platelets were excluded. In this step, the gene annotation “genes shared with humans” (see **Gene annotation and ortholog information**) for each animal species was used. Finally, regarding genes/cell and counts/cell values, cells with >3 |Z score| were excluded.

#### Data integration, visualization, and cell clustering

Data integration, visualization, and cell clustering for each animal species were performed using the Seurat package. In these processes, the expression levels of HSV-1 and SeV were not used. Data integration is a method merging the gene expression count matrices obtained from different experimental conditions while removing batch effects. We integrated the count matrices from the four different conditions for each animal species. In the data integration, SCTransform (a modeling framework for the normalization and variance stabilization of molecular count data from scRNA-seq data) was performed using the SCTransform function for each count matrix. Next, to extract 2000 genes with higher variance and thus greater information for integration, the four count matrices were processed using the SelectIntegrationFeatures function. Next, we used the PrepSCTIntegration function to transform normalized counts into counts per 10,000 counts in the cell (CP10k). After that, we used the FindIntegrationAnchors function with the setting Mock as a reference to find “Integration anchors”. Finally, we integrated the four normalized count matrices using the IntegrateData function with the option ‘normalization.method=“SCT”‘.

For visualization, we first performed principal component analysis (PCA) using the RunPCA function. Then, UMAP^40^ was performed with the RunUMAP function. In this step, principal components (PC) 1-50 were used, and the parameter “n.neighbors” was set individually for each animal species (Hs: 20, Pt: 20, Mm: 50, and Ra: 40).

To define cell clusters in each animal species, we performed graph-based unsupervised clustering (**Fig. S2A**). First, the FindNeighbors function was used, and then, the FindClusters function was used. In these steps, the parameter ‘k.param’ for FindNeghbors was set individually for each animal species (Hs: 12, Pt: 10, Mm: 10, and Ra: 20). The parameter ‘resolution’ for FindClusters was also set individually for each animal species (Hs: 2.0, Pt: 2.2, Mm: 1.7, Ra: 1.2).

#### Cell annotation

Regarding each cluster identified by graph-based unsupervised clustering in the section “**Data integration, visualization, and cell clustering**” (**Fig. S2A**), 11 cell types were manually annotated according to i) the predicted cell type by Azimuth (**Fig. S2B**), ii) the distances between each cluster (**Fig. S2C**), and iii) the correspondence of clusters between animal species (**Fig. S2D–F**). First, reference-based cell type prediction was performed using Azimuth for the mock data from each animal species (**Fig. S2B**). In this step, the gene annotation “genes shared with humans” (see **Gene annotation and ortholog information**) for each animal species was used. We checked the enrichment of each predicted cell type in each cluster by Azimuth. Second, we checked the similarities between clusters by hierarchical clustering (**Fig. S2C**) using the mean values of PCs 1-50 among the individual cells (see **Data integration, visualization, and cell clustering**) in each cluster. Notably, PCA was performed using the expression levels of “all genes” (see **Gene annotation and ortholog information**). The Euclidian distance was used for clustering by Ward’s method. Third, to check the correspondence between clusters in each animal species, we performed data integration, clustering, and visualization for mock data from all four animal species (**Fig. S2D–F**). In the integration, the mock data from humans were used as reference data. In this step, the gene annotation “common genes” (see **Gene annotation and ortholog information**) was used.

After categorizing cells into 11 cell types, the 11 cell types were coarse-grained into 6 cell types based on the results of hierarchical clustering analysis (see **Hierarchical clustering**). The six cell types were used in the subsequent analysis.

### Quantification and Statistical Analysis

#### Hierarchical clustering

To examine the similarities in expression patterns among the conditions (4 animal species × 4 stimuli × 6 cell types = 96 conditions), hierarchical clustering analysis was performed. In this analysis, the 5,000 genes with the highest median absolute deviation (mad) values were used (**Fig. S2H**). First, the average expression levels of the respective genes in each condition were calculated. Next, PCA was performed using the average expression profiles. Third, using PCs 1-30, the distance matrix for the 96 conditions was generated using 1−Pearson’s correlation coefficient. Finally, hierarchical clustering by Ward’s method was performed using the distance matrix.

To determine which factor (e.g., animal species, stimulus, or cell type) was the most impactful on the gene expression in immune cells, hierarchical clustering was performed using induction patterns upon stimulation (**Fig. 1E**).

Unlike for the results shown in **Fig. S2H**, FC values were used to perform PCA. This analysis used 7557 genes, the union of the top 6000 genes related to total expression levels in the expression profiles of each animal species. The FC expression values (stimulated vs. unstimulated conditions) of those genes were calculated for each cell type in each animal species. To avoid generating infinite FC values, the data for genes with zero expression in mock data were set at the minimum nonzero expression level in the mock data. Finally, hierarchical clustering was performed using the method described above.

#### Tensor decomposition

To extract species-specific/common induction patterns upon stimulation from transcriptome data with complex structures (4 animal species × 3 stimuli × 4 cell types × 7557 orthologous genes), we used tensor decomposition (**Fig. 2A**). As the input data for tensor decomposition, the FC values of 7557 genes, the union of the top 6000 genes related to total expression levels in the expression profiles of each animal, were used. The calculation method for FC values is described in the section “**Hierarchical clustering**”. The standardized FC values for each condition were represented as a 4-mode tensor (animal species × stimulus × cell type × orthologous gene). To perform Tucker decomposition (TD), a method of tensor decomposition, we used TensorLy (v0.6.0) (http://tensorly.org/stable/index.html). We performed TD via higher-order orthogonal iteration (HOI) with the parameter ‘init=“svd”‘. In HOI, the size of the core tensor (ranks) was set as [animal species: 3, stimulus: 2, cell type: 3, gene: 15]. The number of iterations was set as 100.

#### Gene classification using the tensor decomposition results

A schematic of the gene classification using tensor decomposition is shown in **Fig. 2C and Fig. S3C–F**. Briefly, we selected the candidate gene categories that had patterns of values (high, mid, or low) (**Fig. S3C**) that matched the ideal pattern (**Fig. S3D**) and then selected the gene category with the best “similarity score” (**Fig. S3E**) from the candidates as the gene category for that gene (**Fig. S3F**).

Initially, the product of the core tensor and the three factor-matrices, A2 (for stimulus), A3 (for cell type), and A4 (for gene), was calculated to obtain three cubic data with three axes, stimulus, cell type, and gene, using the ttl function of rTensor (v1.4.8) (https://github.com/rikenbit/rTensor). Each cubic data point indicated information related to species-common, bat-specific, and macaque-specific factors (**Fig. 2B**). Next, since the values of latent factors related to HSV-1 and SeV were similar (**Fig. S3A**), these two categories were integrated into the category “Virus” by calculating mean values. Additionally, since the values of latent factors related to NaiveT and KillerTNK were similar (**Fig. S3B**), these two categories of cell types were integrated into the category “TNK” by calculating mean values. Thus, hereafter, the category of stimuli included virus and LPS, and the category of cell types included B cells, TNK cells and Monos.

Then, in each cubic data, genes were classified into 11 categories (**Fig. 2C**) through the following three steps. Briefly, from the candidate gene categories that had patterns of values (high, mid, or low) (**Fig. S3C**) that matched the ideal pattern (**Fig. S3D**), the gene category with the lowest “similarity score” (**Fig. S3E**) was selected as the gene category for that gene (**Fig. S3F**).

In the first step (**Fig. S3C**), the values in each cubic data were normalized, and the genes were classified into three classes (high, mid, and low) according to the ranking of values in each condition (stimulus × cell type). First, six column vectors in the TD results for the 6 conditions (2 stimuli × 3 cell types) were normalized by dividing them by the 90th percentile for the individual vectors. After the division step, to suppress the effect of abnormally high or low values, data with > 1 or < -1 were assigned as 1 and -1, respectively. Next, the genes were categorized into three classes based on the rule that if the rank of a value was greater than the 80th percentile or smaller than the 20th percentile, it was categorized as “high” or “low”, respectively; otherwise, it was categorized as “mid”.

In the second step (**Fig. S3E**), a “similarity score” was calculated to represent the similarity between the genewise pattern of the TD results and the “ideal patterns” for each gene category. The “ideal patterns” were defined as vectors composed of 1, 0, and -1 for 16 gene categories (Virus_high, LPS_low, Virus_low, LPS_high, B_high, TNKM_low, B_low, TNKM_high, TNK_high, BM_low, TNK_low, BM_high, M_high, BTNK_low, M_low, and BTNK_high) (**Fig. S3D**). The “similarity score” was defined as the sum of the residual squares between the two vectors, the genewise vector of normalized values from the TD results (**Fig. S3C**) and the “ideal patterns” (**Fig. S3D**). According to the definition, the “similarity scores” for every combination of genes and gene categories were calculated. After calculating all similarity scores, to obtain the threshold for checking if a gene should be recognized as a gene in that category, the 20th percentile of the similarity score in the vector for each gene category was calculated.

In the third step (**Fig. S3F**), the gene category for each gene was determined. First, the candidate gene categories for each gene were filtered according to the pattern assigned in the first step (**Fig. S3C**). If the pattern (high/mid/low) of all 6 conditions was high or low, the gene was categorized as ALL_high or ALL_low, respectively. If the pattern of a gene matched the “ideal pattern” of a gene category, the gene category was added as a candidate gene category for the gene. For example, if the pattern of gene A was (Virus_B: high, Virus_TNK: high, Virus_M: high, LPS_B: high, LPS_TNK: low, LPS_M: mid), the candidate gene category for gene A was “Virus_high” and “B_high” because all virus-infected data were assigned as “high” and all B-cell data were assigned as “high” (**Fig. S3D**). Second, the gene category with the lowest “similarity score” among the candidate gene categories was selected as the tentative gene category. In this selection, if the “similarity score” was higher than the threshold of the gene category (**Fig. S3E**), the gene was categorized as “Others” (See gene B in **Fig. S3F**) because the pattern for the gene was recognized as being too different from the “ideal pattern”. If no candidate gene category was available, the gene was also classified as “Others” (See gene C in **Fig. S3F**). Finally, the final gene category was determined by integrating similar gene categories (**Fig. S3F**). For instance, the categories Virus_high and LPS_low were integrated into the category Virus_high because both categories indicated that virus-infected data were higher than LPS-stimulated data (See gene D in **Fig. S3F**). As a result of the gene classification process, genes were categorized into one of 11 categories (**Fig. 2C, S3D**).

#### GO term enrichment analysis

Gene Ontology (GO) analysis was performed with Fisher’s exact test. This analysis used the GO canonical pathways and GO biological processes defined by MSigDB (v7.3) (https://www.gsea-msigdb.org/gsea/msigdb/collections.jsp). Adjusted P values were calculated using the Benjamini□Hochberg (BH) method.

#### Calculation of gene set variation analysis (GSVA) scores

The gene set-wise expression scores used in **Figs. 3A, 4B, 4C, and S4B** were calculated using GSVA (v1.38.2)^41^ with the algorithm “ssgsea”.

#### Identification of differentially expressed genes (DEGs) and marker genes

In bat monocytes, DEGs were identified in Cluster 5 or Cluster 7 compared to the other clusters using the FindMarkers function of Seurat packages. A gene that met the following three criteria was considered a DEG: 1) the false discovery rate (FDR) calculated using the BH method was less than 0.05, 2) the average log2FC was greater than 1 or less than -1, and 3) the proportion of expressing cells was greater than 0.2.

The marker genes of Cluster 5 and Cluster 7 of bat monocytes (RaC5marker and RaC7marker, respectively) were defined as upregulated DEGs in Cluster 5 (**Fig. 4E**) and Cluster 7 (**Fig. 4G**), respectively.

## [Supplemental Information]

**Figure S1. Validation of viral infectivity and the innate immune response (related to Figure 1)**

(A) Heatmap of the induction levels of genes related to the IFN response and inflammation. The rows indicate genes, and the columns indicate combinations of species, stimulus, and dose. The color represents the log2 Fold Change of ddCt upon stimulation measured by qRT□PCR. “rep. 1” and “rep. 2” indicate biological replicates.

(B-C) Heatmap of the expression levels of viral genes (B: HSV-1; C: SeV) measured by qRT□PCR. The rows indicate viral genes, and the columns indicate combinations of species and doses. “rep. 1” and “rep. 2” indicate biological replicates. The color represents the ddCt values based on the expression levels of GAPDH.

(D-G) Violin plots of (D) the numbers of detected genes per cell before QC, (E) numbers of counted reads per cell before QC, (F) numbers of detected genes per cell after QC, and (G) numbers of counted reads per cell after QC.

**Figure S2. Heterogeneous expression patterns in the four animal species (related to Figure 1)**

(A-B) UMAP plots representing the gene expression patterns of PBMCs from the four species. Each dot is colored according to the results of unsupervised clustering (A) and reference-based label transfer (B).

(C) Heatmaps showing pairwise Euclid distances representing the gene expression differences among clusters. The distances were calculated using PCs 1-50 of the gene expression data.

(D-E) UMAP plots representing the gene expression patterns of PBMCs from the mock samples for the four species. Each dot is colored according to the results of unsupervised clustering using the integrated data for the four mock samples (D) or the four samples from each animal shown in **Figure S2A** (E).

(F) Heatmaps showing pairwise Euclid distances representing the gene expression differences among clusters shown in **Figure S2D**. The distances were calculated using PCs 1-30 of the gene expression data.

(G) Dot plots representing the expression patterns of marker genes for each cell type defined by Azimuth (azimuth.hubmapconsortium.org/references/#Human%20-%20PBMC)

(H) Hierarchical clustering analysis of 48 pseudobulked FC gene expression datapoints (4 animal species x 4 stimuli x 11 cell types = 176 conditions).

**Figure S3. Classification of genes according to species-specific expression patterns (related to Figure 2)**

(A) Heatmap representing a latent factor matrix related to stimuli. The columns indicate stimuli, and the rows indicate latent factors representing stimulus-common (L2_1) and virus vs. LPS (L2_2) factors.

(B) Heatmap representing a latent factor matrix related to cell types. The columns indicate cell types, and the rows indicate latent factors representing cell type-common (L3_1), monocyte-specific (L3_2), and B-cell-specific (L1_3) factors.

(C) Summary of the normalization of values and patterning according to the ranking of the values. First, six column vectors (2 stimuli × 3 cell types) in the TD results were normalized by dividing them by the 90th percentile of the individual vectors. Then, data with > 1 or < -1 were assigned as 1 and -1, respectively. Next, the genes were categorized into three classes (high, mid, and low) based on the rule that if the rank of a value was greater than the 80th percentile or smaller than the 20th percentile, it was categorized as “high” or “low”, respectively; otherwise, it was categorized as “mid”.

(D) Summary of the ideal patterns for each gene category used in the gene classification in **Figure 2C**.

(E) Summary of the calculation of the similarity score and establishment of the threshold for the gene classification in **Fig. S3F**. The sum of the residual squares between two vectors, the genewise vector of normalized values from the TD results (**Fig. S3C**) and the “ideal patterns” (**Fig. S3D**) were calculated. Then, the threshold used in **Fig. S3F** was obtained by calculating the 20th percentile of the similarity score for the vector for each gene category.

(F) Summary of gene classification. By comparing patterns from the TD results (**Fig. S3C**) and the ideal patterns (**Fig. S3D**), candidate gene categories were selected. Next, the gene category with the lowest “similarity score” among the candidate gene categories was selected as the tentative gene category. In this selection, if the “similarity score” was higher than the threshold of the gene category (**Fig. S3E**), the gene was categorized as “Others” (gene B). If no candidate gene category was available, the gene was also classified as “Others” (gene C). Finally, the final gene category was determined by integrating similar gene categories (genes A and D).

(G-I) Heatmap representing the values of the products calculated in **Figure 2C**. The data relating to (G) the species-common factor (L1_1), (H) the bat-specific factor (L1_2), and (I) the macaque-specific factor (L1_3) are shown. Each row indicates the respective gene. The color keys shown on the right of the heatmap indicate gene categories.

(J-L) Heatmap representing the FC values in the input tensor. The orders of the rows are the same as in (J) **Figure S3G**, (K) **Figure S3H**, and (L) **Figure S3I**. Each row indicates the respective gene. The color keys shown on the right of the heatmap indicate gene categories.

**Figure S4. Identification of species-specific cell types (related to Figure 4)**

(A) UMAP plots representing the expression patterns of every single cell. Dimensionality reduction was performed for each combination of the four species and three cell types.

(B) UMAP plots representing the average expression levels of marker genes for Cluster 7 [C7markers].

**Table. S1. Primers used for RT □**_**qPCR (related to the STAR Methods)**

The sequences of the primers used for RT□qPCR are listed.

