## Supplementary figures and images for "Single-cell transcriptome analysis illuminating the characteristics of species-specific innate immune responses against viral infections"

### Figure S1

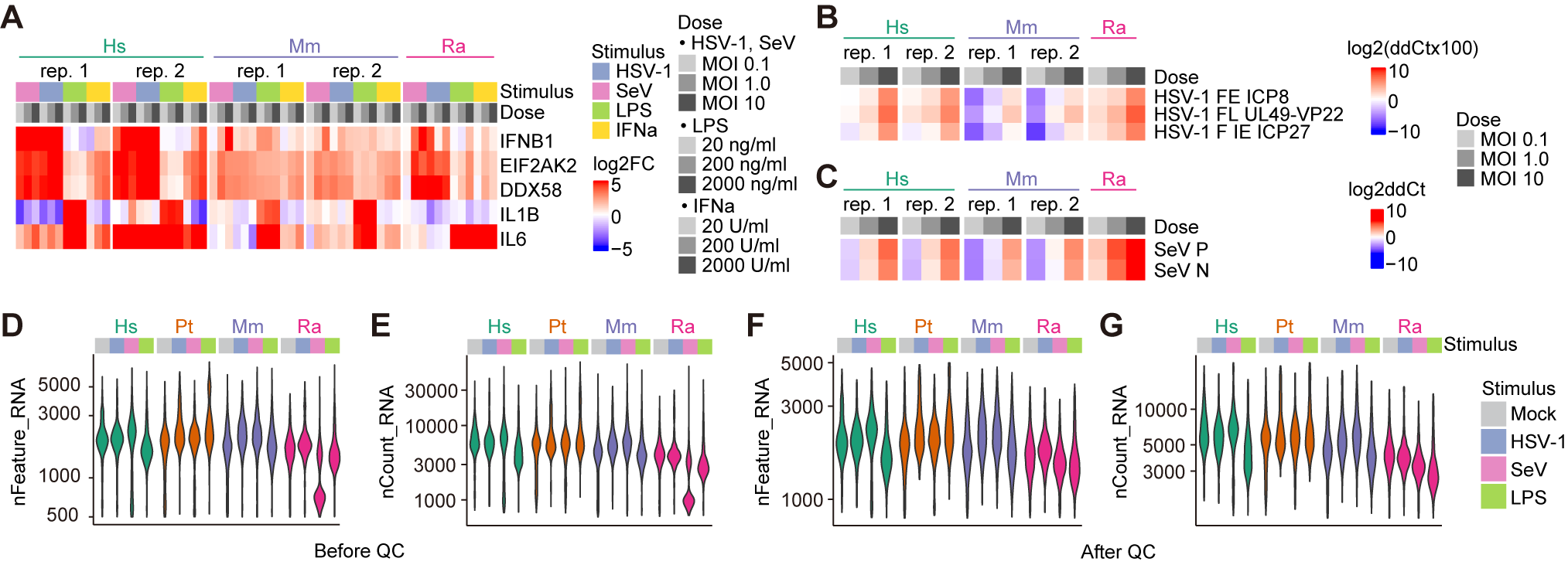

### Figure S2

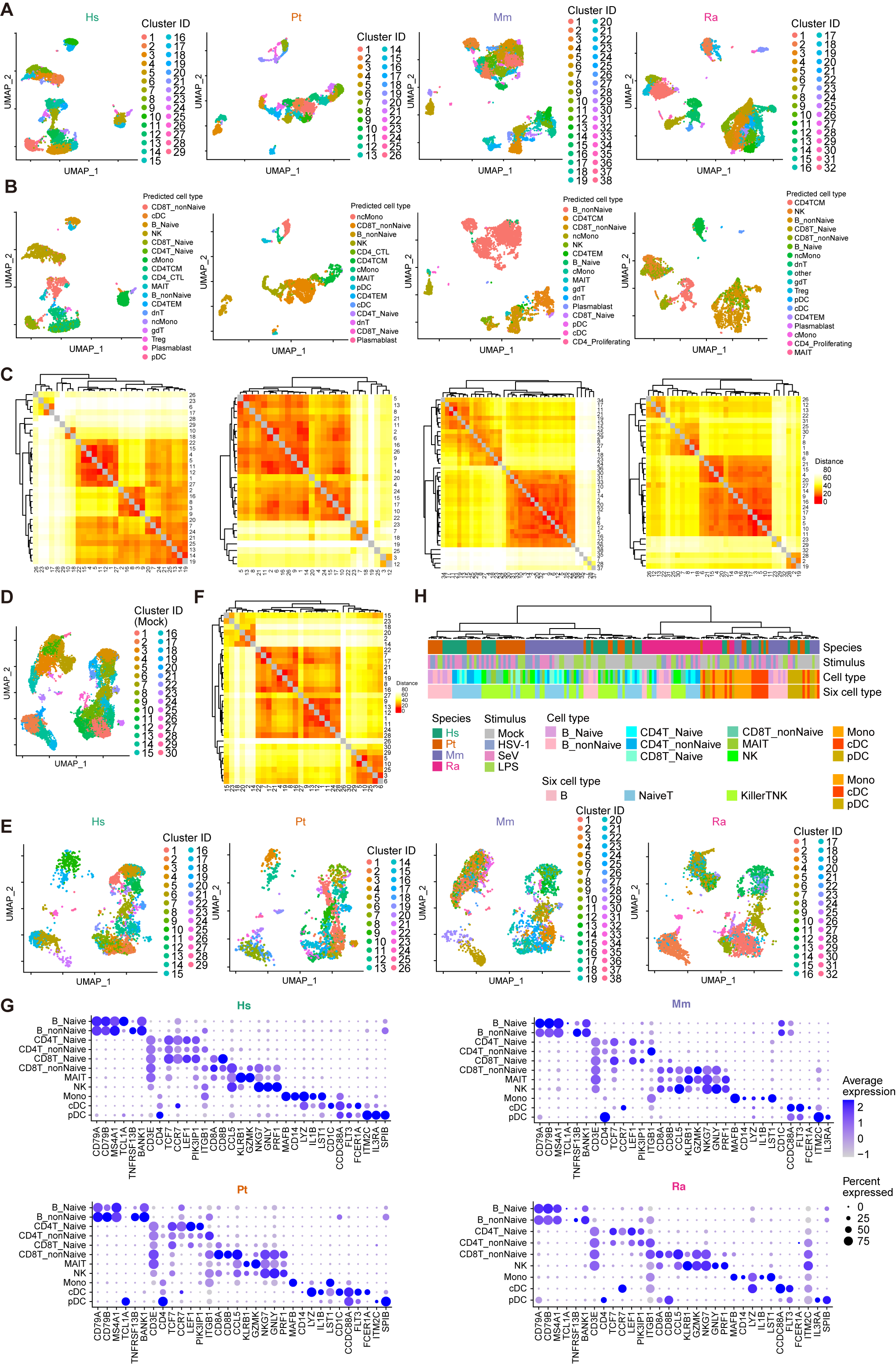

### Figure S3

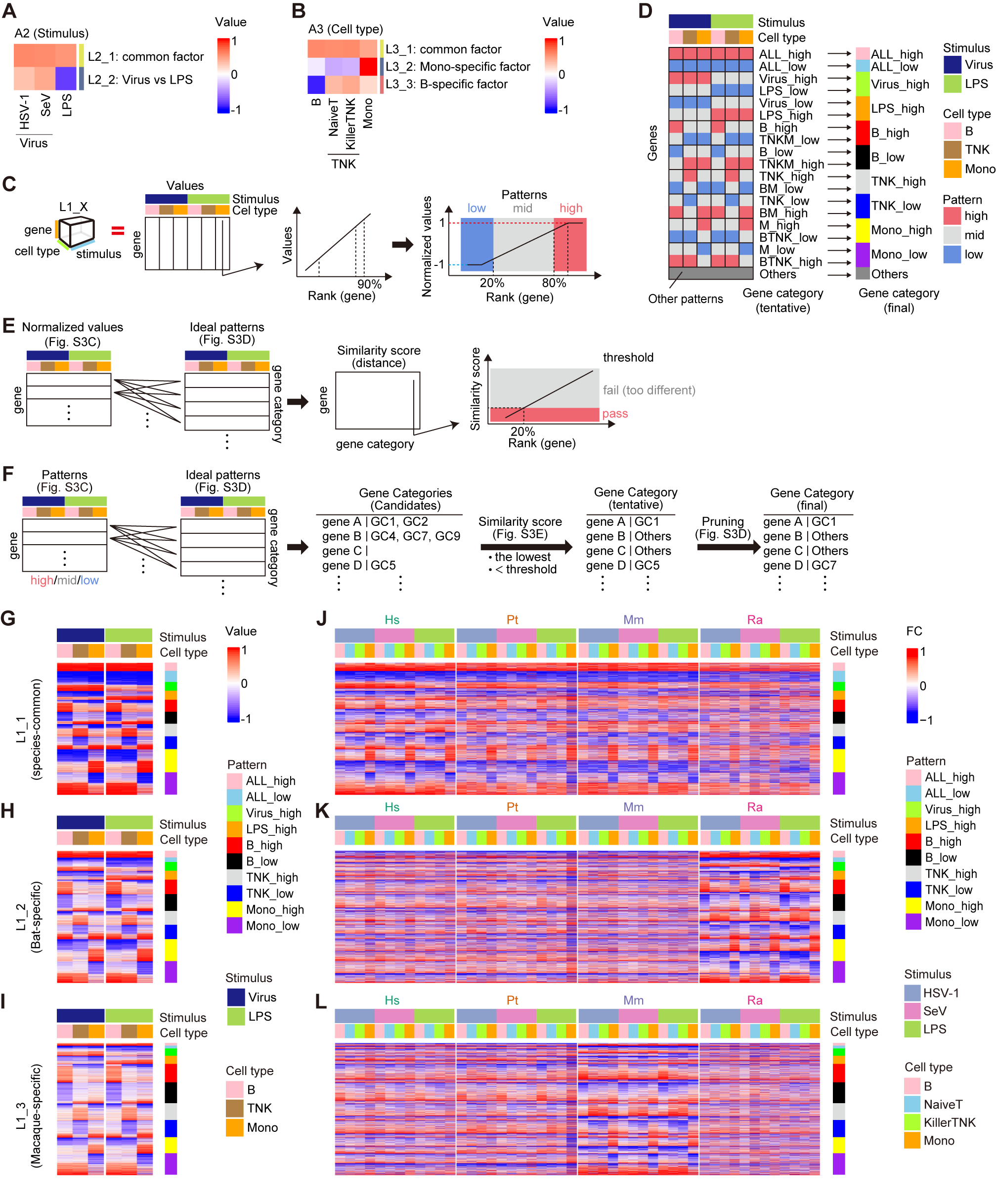

### Figure S4

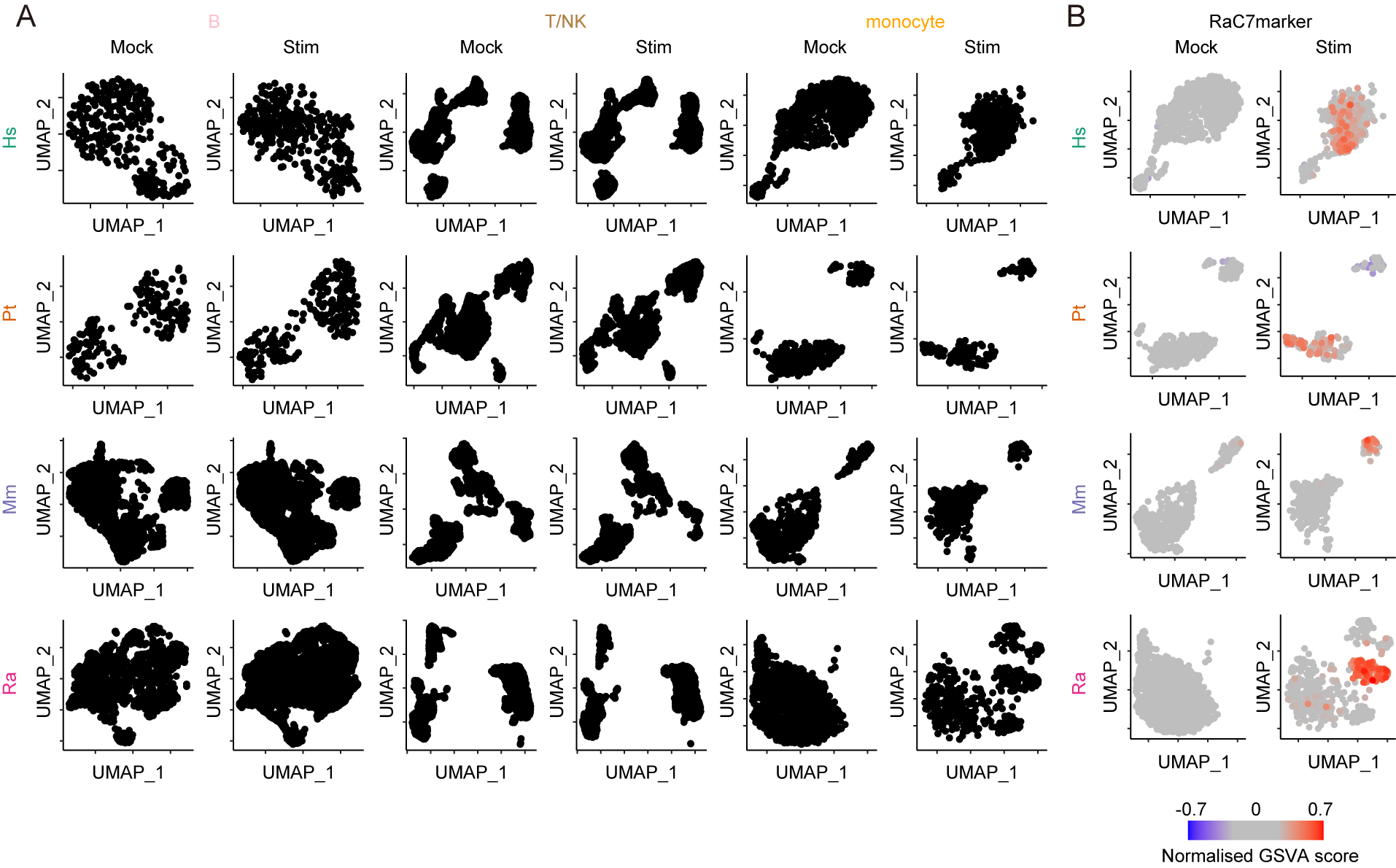
